# Blind testing cross-linking/mass spectrometry under the auspices of the 11^th^ critical assessment of methods of protein structure prediction (CASP11)

**DOI:** 10.1101/053173

**Authors:** Adam Belsom, Michael Schneider, Lutz Fischer, Oliver Brock, Juri Rappsilber

## Abstract

Determining the structure of a protein by any method requires varies contributions from experimental and computational sides. In a recent study, high-density cross-linking/mass spectrometry data in combination with *ab initio* structure prediction by conformational space search determined the structure of human serum albumin (HSA) domains, with an RMSD to X-ray structure of up to 2.53 Å, or 3.38 Å in the context of blood serum. This paper reports the blind test on the readiness of this technology through the help of Critical Assessment of protein Structure Prediction (CASP). We identified between 201-381 unique residue pairs at an estimated 5% FDR (at link level albeit with missing site assignment precision evaluation), for the four proteins that we provided data for. This equates to between 0.63-1.20 proximal residues per residue, which is comparable to that obtained in the HSA study (0.85 links per residue at 5% FDR). Nevertheless, initial results of CASP11 have suggested that improvements in structure prediction using cross-link data are slight. Most significantly, however, CASP11 revealed to us some of the current limitations of cross-linking, spelling out areas in which the method must develop in future: links spread unevenly over sequence and beta sheets both lacked links and suffered from weak definition of observed links over structure. With CASP12 taking place this year and biannually in the future, blind testing low-resolution structure analysis tools is a worthwhile and feasible undertaking. Data are available via ProteomeXchange with identifier PXD003643.

**The abbreviations used are:**

CLMS: cross-linking/mass spectrometry;
NHS: *N-*hydroxysuccinimide;
NMR: nuclear magnetic resonance;
sulfo-SDA: sulfo-NHSdiazirine, sulfosuccinimidyl 4,4’-azipentanoate;
FDR: false discovery rate;
MBS: model-based search;
HSA: human serum albumin;
RMSD: root-mean-square deviation;
CASP: Critical Assessment of protein Structure Prediction;
Tris: tris(hydroxymethyl)aminomethane;
PES: polyethersulphone;
IAA: iodoacetamide;
LTQ: linear trap quadrupole;
MS2: tandem MS scan;
LC-MS: liquid chromatography mass spectrometry;
FM: free modelling.

## INTRODUCTION

Cross-linking/mass spectrometry (CLMS) is a well-established, low-resolution technique for revealing protein interactions in protein complexes and studying protein conformational changes (1–4). CLMS successes to date are numerous for multi-protein complexes (5) and include the production of a topological map of the Nup84 complex (6), insights on the structure of the NDC80 complex (7), determination of the structure of the RNA polymerase II-TFIIF complex (8), ribosomal protein S1 (9), nucleoporin Nup133 (10), coatomer-related heptameric molecule from the nuclear pore complex (11), GTP-bound translation initiation factor 2, eIF2B (12) the structure of the mammalian mitochondrial ribosome (13, 14), and most recently, revealing dynamic aspects regarding the maturation of the proteasome lid complex (15), or validating insights on the structure of isolated complexes also in their native environments (16).

In contrast to these CLMS successes, the use of cross-linking data in resolving a detailed protein structure is less well established, of which a critical obstacle has been the low data density obtainable using the standard NHS-ester based cross-linking chemistry. We found 0.07 constraints per residue in earlier work from our lab (17) and a study attempting to increase linked residue pair numbers for 8 model proteins, using size exclusion chromatography and multiple proteases, identified between 0.01-0.09 constraints per residue (18). Young *et al*. however, managed to identify the correct fold of bovine basic fibroblast growth factor by combining such low-density cross-linking data with threading and homology modelling (19). We believe that CLMS holds the promise of becoming an approach to protein structure determination that complements established methods including nuclear magnetic resonance (NMR) spectroscopy, X-ray and cryo-electron-microscopy. The number of cross-links per amino acid residue required for protein structure prediction is currently unknown, but we might reasonably expect the value to be close to the 3-20 constraints per residue obtained in NMR spectroscopy (20).

We have worked on improving CLMS to significantly increase the density of cross-links per protein. Most prior CLMS studies have relied on the DSS/BS3 cross-linker, involving homobifunctional NHS-ester chemistry, predominantly involving lysine side chains. As this limits cross-linking sites to lysines, the resulting linkage maps lack the data density necessary to define the detailed structure of proteins. In principle, the use of cross-linkers based on photoactivatable diazirine should widen the range of cross-linkable amino acids (21–27). Indeed, use of sulfo-NHS-diazirine (sulfo-SDA) allowed the number of cross-links observed in a protein to be greatly increased, by increasing the number of reactive amino acids (28) from 4 x 4 (K, S, T and Y residues only) to 4 x 20 (K, S, T and Y to any other residue), opening up the possibility of determining entire protein structures. However, the high promiscuity of the linker also exacerbates the already established need for false discovery rate (FDR) estimation (7, 29) and drives the requirement for further developments on estimating the rate of falsely called residue pairs (Fischer and Rappsilber, submitted). In previous work, we showed that combining the resulting high-density photo-cross-linking/mass spectrometry (photo-CLMS) data with the conformational space search algorithm model-based search (MBS) (30) can produce accurate protein structure models of the domains of human serum albumin (HSA) in purified form (RMSD to crystal structure of 2.53 Å) and in its native environment, blood serum (RMSD to crystal structure 3.38 Å) (28).

Solving domain structures of HSA provided the proof-of-concept for photo-CLMS in tandem with computational methods as a protein structure determination technology. We found that having 2.5 constraints per amino acid residue was sufficient to determine these domain structures with RMSD to crystal structure accuracies of 2.5/4.9/2.9 Å (28). Yet it left open which experimental details are critical to this success and also how widely applicable this technology would be at the current time. We therefore embarked on a blind study to evaluate the current capabilities of photo-CLMS with regards to structure determination and teamed up with the organizers (in particular Krzyzstof Fidelis and Andriy Kryshtafovych) of Critical Assessment of protein Structure Prediction (CASP).

CASP is a community-wide experiment for protein structure prediction taking place every two years since 1994 (31, 32). CASP provides the opportunity for the protein modeling community to test their computational algorithms and, consequentially, is also a test of the state of the art in structure prediction (33). The experiment depends on “blind prediction”, that is, at the time of modeling the experimental structure has not yet been published and is unknown to the modelers. The CASP Organizing Committee obtains early access to solved but withheld protein structures and then decides upon which of the proteins are suitable for the test and subsequently releases the protein sequences to all participants. Proteins are typically interesting from a biological perspective, can involve unique protein folds and exhibit little homology with known protein structure. Therefore, the protein targets in CASP are typically challenging. Models are then submitted to the CASP server and evaluated by a team of independent assessors after the submission deadline (34). It is not unusual for modelers to fail to correctly predict a structurally accurate model; learning from the failures however, allows improvements to be made to the algorithms (35).

In discussion with the CASP Organizing Committee, we agreed to apply our experimental pipeline for CLMS to the blind study of potentially challenging proteins for CASP11. The Rappsilber lab received protein sequences and the actual proteins. After putting the proteins through the CLMS pipeline, the Rappsilber lab then submitted CLMS data in the form of distance restraints to a CASP server, prior to release of the protein structure to the prediction community. Predictors were then able to include the CLMS data into their predictions.

We saw the CASP experiment as an excellent opportunity to perform a rigorous blind test of our CLMS method. We hypothesized that such a test would play an important role in assessing the current capabilities of this technology and in suggesting improvements to increase method accuracy and robustness. Our data also hold tremendous promise for the modeling field, which is analyzed and discussed elsewhere (36).

## EXPERIMENTAL PROCEDURES

### Proteins

A total of nine proteins were received from five labs. YaaA was received from the lab of Prof. Mark Wilson (Department of Biochemistry/Redox Biology, University of Nebraska), as a frozen solution (25 mM HEPES, pH 8.2, 100 mM KCl, 6.89 mg/mL). Five proteins: 413472 (GS13694A), BACUNI_01052, RUMGNA_02398, SAV1486 and BACCAC_02064 were received from the lab of Dr. Ashley M. Deacon (Joint Center for Structural Genomics (JCSG), Stanford Synchrotron Radiation Lightsource, Stanford University). All were received as previously frozen and thawed-in-transit solutions, with all comprised of a buffer containing 20 mM tris(hydroxymethyl)aminomethane (Tris), pH 7.9, 150 mM NaCl, 0.5 mM tris(2-carboxethyl)phosphine (TCEP) and at concentrations of 2.3, 4.6, 5.2, 2.5 and 11 mg/mL respectively. MmR495A was received from the lab of Prof. Gaetano Montelione (Center for Advanced Biotechnology and Medicine, Rutgers University), as both a solid lyophilisate (from 20 mM NH4OAc) that had absorbed water during transit and also as a frozen solution on ice containing 10 mM Tris, pH 7.5, 100 mM NaCl, 10 mM DTT, 0.02% NaN_3_. Af1502 was received from the lab of Dr. Jörg Martin (Max-Planck Institute for Developmental Biology, Tübingen) as a frozen solution of 30 mM MOPS, 250 mM NaCl, 10% glycerol, pH 7.2, 16 mg/mL. Laminin was received from the lab of Prof. Deborah Fass (Department of Structural Biology, Weizmann Institute of Science) as a frozen solution on ice containing PBS and 10% glycerol, 2.4 mg/mL.

Four of the six designated targets (BACUNI_01052, RUMGNA_02398, SAV1486 and BACCAC_02064) were buffer-exchanged prior to cross-linking to remove Tris from the buffer. Buffer exchange was carried out using polyethersulphone (PES) ultracentrifugation devices for concentration of small-volume protein samples, Vivaspin 500, 5000 MWCO, GE Healthcare. Protein concentration was estimated using a Nanodrop 1000 Spectrophotometer from Thermo Fisher Scientific, measuring at 280 nm.

### Chemical Cross-linking

Each target was cross-linked using sulfo-SDA, using four different cross-linker to protein ratios (2:1, 1:1, 0.5:1 and 0.25:1, w/w) and four UV activation times (15, 30, 45 and 60 minutes). Cross-linking was carried out in two-stages: firstly sulfo-SDA dissolved in cross-linking buffer (20 mM HEPES-OH, 20 mM NaCl, 5 mM MgCl_2_, pH 7.8) was added to target protein (30 μg, 1 μg/μL) and left to react in the dark for 1h at room temperature. This allowed the reaction of lysine side chain amino groups but also hydroxyl groups in serine, threonine and tyrosine side chains, with the sulfo-NHS ester component of the cross-linker. The diazirine group was then photo-activated using UV irradiation, at 365 nm, from a UVP CL-1000 UV Crosslinker (UVP Inc.). Samples were spread onto the inside of Eppendorf tube lids by pipetting (covering the entire surface of the inner lid), placed on ice at a distance of 5 cm from the tubes and irradiated for either 15, 30, 45 or 60 minutes. Following the reaction, half of each reaction condition sample was then pooled as a “mixed” sample (a total of 240 μg). The resulting cross-linked mixtures were then separated by electrophoresis using a NuPAGE 4-12% Bis-Tris gel, ran using MES running buffer and stained using Imperial Protein Stain from Thermo Scientific, a Coomassie blue stain. Protein monomer bands were excised from the gel, cut into pieces and then washed to remove Coomassie staining. Proteins were reduced with 20 mM DTT, alkylated using 55 mM IAA and digested overnight using trypsin following standard protocols (7). Trypsin/Glu-C co-digestion (in-gel trypsin digestion, overnight at 37 °C followed by addition of Glu-C for 6 hours at room temperature) was used for mixed samples of Targets 1 and 2. In addition, in-solution Glu-C digestion was used for mixed samples of Targets 2-4. Digests were desalted using self-made C18 StageTips (37) prior to mass spectrometric analysis.

### Mass Spectrometry and Data Analysis

Peptides were loaded directly onto a spray emitter analytical column (75 μm inner diameter, 8 μm opening, 250 mm length; New Objectives) packed with C18 material (ReproSil-Pur C18-AQ 3 μm; Dr Maisch GmbH, Ammerbuch-Entringen, Germany) using an air pressure pump (Proxeon Biosystems) (38). Mobile phase A consisted of water and 0.1% formic acid. Mobile phase B consisted of acetonitrile and 0.1% formic acid. Peptides were loaded onto the column with 1% B at 700 nl/min flow rate and eluted at 300 nl/min flow rate with a gradient: 1 minute linear increase from 1% B to 9% B; linear increase to 35% B in 169 minutes; 5 minutes increase to 85% B. Eluted peptides were sprayed directly into a hybrid linear ion trap - Orbitrap mass spectrometer (LTQ-Orbitrap Velos, Thermo Fisher Scientific). Peptides were analyzed using a “high/high” acquisition strategy, detecting peptides at high resolution in the Orbitrap and analyzing, also in the Orbitrap, the products of their CID fragmentation in the ion trap. Survey scan (MS) spectra were recorded in the Orbitrap at 100,000 resolution. The eight most intense signals in the survey scan for each acquisition cycle were isolated with an *m/z* window of 2 Th and fragmented with collision-induced dissociation (CID) in the ion trap. 1* and 2* ions were excluded from fragmentation. Fragmentation (MS2) spectra were acquired in the Orbitrap at 7500 resolution. Dynamic exclusion was enabled with 90 seconds exclusion time and repeat count equal to 1.

### Data analysis

Mass spectrometric raw files were processed into peak lists using MaxQuant version 1.3.0.5 (39) using default parameters except the setting for “Top MS/MS peaks per 100 Da” being set to 100. Peak lists were searched against a database, comprising in each case only the sequence of the protein that was being analyzed using Xi (ERI, Edinburgh) for identification of cross-linked peptides. Sequences were provided by the CASP Organizing Committee and are available in the supplement. Search parameters were MS accuracy, 6 ppm; MS/MS accuracy, 20 ppm; enzyme, trypsin; specificity, fully tryptic; allowed number of missed cleavages, four; cross-linker, SDA; fixed modifications, none; variable modifications, carbamidomethylation on cysteine, oxidation on methionine, SDA-loop (SDA cross-link within a peptide that is also cross-linked to a separate peptide). Other SDA modifications (including those resulting from reaction with water and ammonia) were not included in the database search, as in earlier work we identified very few such modifications and including these modifications served to increase search database size and also increase false positive identifications, which we were keen to avoid in this experiment. Linkage specificity for sulfo-SDA was assumed to be at lysine, serine, threonine, tyrosine and protein N-termini at one end, with the other end having no specificity, i.e. linking to any amino acid residue. A modified target-decoy search strategy was used to estimate FDR (7, 29) [Fischer and Rappsilber, submitted]. In short, unique residue pairs are scored on supporting PSMs by,

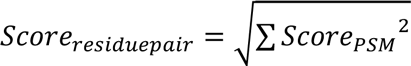

Before scoring residue pairs and applying an FDR at their level, the dataset is pre-filtered by applying an FDR-based score cut-off for PSMs and a subsequent FDR-based score cut-off for unique peptide pairs (scored the same way as residue pairs based on supporting PSMs). This provides a means to do noise filtering and can increase the number of unique residue pairs that pass a given FDR. Optimal score cut-offs are automatically defined using xiFDR: https://github.com/lutzfischer/xiFDR.

The MS data have been deposited to the ProteomeXchange Consortium via the PRIDE partner repository with the dataset identifier PXD003643 (40) (Reviewer account details: Username, Password: ipIq98i4).

## RESULTS AND DISCUSSION

### Experiment initial setup

The pairing of our lab with CASP11 tested the current capabilities of CLMS in the context of a specific set of proteins, selected for being high-level protein structure prediction targets. In addition, CASP participants tested the current value of such experimental data for protein structure prediction. The latter aspect is discussed in a separate manuscript by a participating modeling lab (36). Please note that throughout the entire CASP experiment, the structure of the protein targets was unknown to our group and to the modeling groups.

Structural genomics centers, who are in collaboration with the CASP Organizing Committee, had agreed to withhold the release of their newly resolved structures to the Protein Data Bank (PDB) (Fig. 1a). This allowed the blind nature of the experiment to be maintained. The CASP Organizing Committee assessed the suitability of protein structures as potential targets for the CASP11 experiment. Structures were considered for different experiments within CASP11, one of which involved CLMS-assisted structure prediction, known within CASP11 as “contact-guided prediction”. Structures selected for the contact-guided prediction experiment were believed to be particularly challenging for structure predictors. Structural genomics centers providing protein structures, then sent quantities of selected target protein to our lab for cross-linking, data acquisition, data processing and FDR estimation. Cross-link data was then sent to the CASP Organizing Committee which released the data on their website to initiate the second part of the experiment, testing current prediction gains achieved by CLMS data. Both elements of the experiment ended with the release of the respective structures via PDB.

**FIG. 1.**
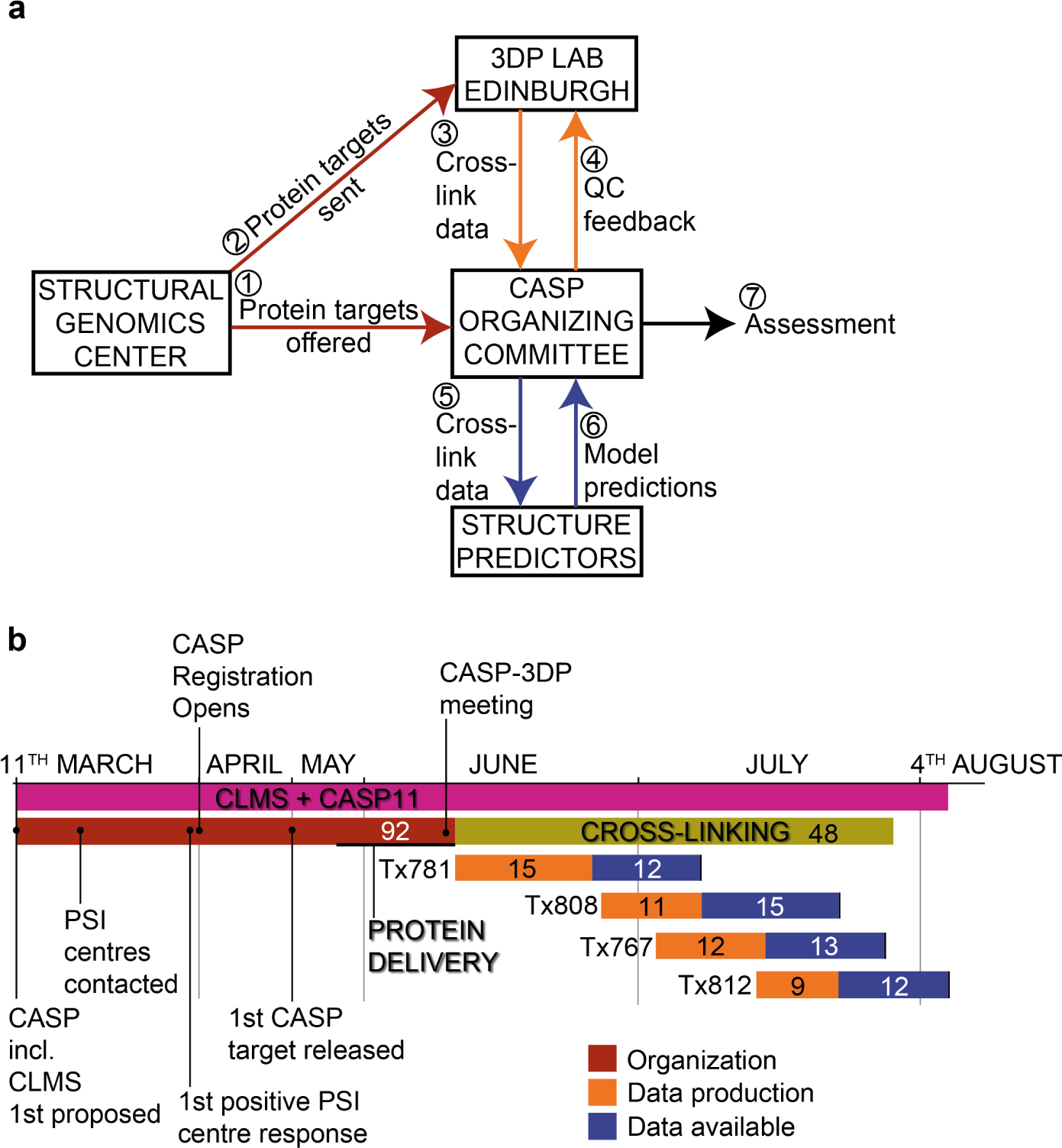
Organisation of CASP11 including cross-linking/mass spectrometry. (a) Schematic representation of the interactions between CASP11 participants. Numbers represent the order in which each interaction took place. (b) CASP11 and CLMS timeline. Total duration of CASP11 and CLMS denoted by *pink bar*. Duration of organisational aspects denoted by *red bar*. Duration of the cross-linking and mass spectrometry aspect denoted by *green bar*. Each of the four targets for which data was provided is represented by a separate bar. Each of these bars is in two sections: The duration of the data production element is represented in *orange* and the duration that data was made available prior to target expiration is represented in *blue*. The numbers within the timeline bars denote the number of days in duration for that particular element.

### Experiment time line

CASP11 including CLMS was first proposed on March 11^th^ 2014 (Fig. 1b). On March 18^th^, the CASP Organizing Committee contacted structure providers to enquire whether proteins could be sent to our lab. The first positive response from a PSI center came 12 days later on March 30^th^. The first proteins arrived to Edinburgh on May 29^th^, with the last proteins arriving on June 9^th^. A meeting was held on June 10^th^ between the CASP Organizing Committee and us to discuss the details concerning the format of the experiment, which of the delivered proteins were to become targets in CASP11 and the order in which data production on the targets was to be done. Cross-linking commenced on June 11^th^. This meant that the organizational phase of the experiment, beginning on March 11^th^, had lasted a total of 92 days. In contrast, protein cross-linking and data processing was carried out for a total of 48 days.

A total of nine proteins (YaaA, 413472 (GS13694A), BACUNI_01052, RUMGNA_02398, SAV1486, BACCAC_02064, laminin, Af1502 and MmR495A) made their way to Edinburgh. One shipment was withheld in customs and arrived defrosted, potentially compromising four targets. Six targets were identified as potential candidates for providing CLMS data. The initial criteria were protein size > 20 kDa, proteins monomeric in solution and having ~1 mg protein sample available (even though we ended up using only 250 μg). We suspected that protein size is an important consideration for CLMS data directed model search, as linked residue pair distance constraints are likely less informative for smaller protein structures, whereas larger proteins are most likely too computationally demanding to model within the time limitations of the experiment. For this reason, Af1502 was discounted as a potential target, having a molecular weight of only 8.3 kDa. The necessity for monomeric proteins was driven by the current requirement that we must avoid cross-linking of homodimeric protein structures, to be certain that observed distance constraints are within a protein structure and not between the monomers of a homodimer. For this reason, MmR495A was discounted as a target. 413472 (GS13694A) was not refined, i.e. no structure was determined and the protein therefore not included in CASP11. A fourth protein to be dismissed, part of the original 6 suitable targets, was YaaA. The protein exhibited total aggregation following cross-linking, according to SDS-PAGE. Given the rigid time limits of the experiment no attempt was made to optimize the conditions in any way and the target was simply dropped.

Five proteins were taken forward. The expiration dates of these targets were staggered, scheduled for July 1^st^, July 8^th^, July 23^rd^, July 28^th^ and August 4^th^, which dictated the order that the targets were tackled. The first target, Tx781, was due to expire on July 1^st^. Analysis started June 11^th^ and ended June 26^th^ with release of the data. This would have left only 5 days for computation by structure predictors (3 working days), however the PDB agreed to delay the release of this target by a further 7 days, hence the final expiry date became July 8^th^. Protein BACUNI_01052, due to be CASP Target Tx771, was originally scheduled also for release on July 8^th^. After work on Tx781, this left no time to cross-link and acquire data on this target, and for this reason the target was dropped from the experiment. Data was provided for four targets: Target 1, RUMGNA_02398, Tx781; Target 2, BACCAC_02064, Tx808; Target 3, SAV1486, Tx767 and Target 4, laminin, Tx812.

### Experiment protocol

Given the time constraints that we were working under, to finish prior to release of the structure, we needed to balance acquisition time with analysis depth. As photo-cross-linking is a very recent development as part of CLMS we do not yet possess optimized protocols and a detailed understanding of all factors that govern yield. In a prior analysis of HSA using sulfo-SDA photoactivatable cross-linking (unpublished data), we observed that the number of unique linked residue pairs identified could be increased using different cross-linker to protein ratios and different UV activation times. As CLMS analysis is a stochastic approach, the number of observed residue pairs increases with repeated LC-MS analysis on identical sample. However the maximum number of observable residue pairs in identical sample plateaus after around 4-5 runs. This observation suggests that the limiting factor for identification of unique linked residue pairs is product dependent and the sample has been analyzed exhaustively, within the limits of our chromatography and mass spectrometer. In other words, if we can increase the number of cross-linked peptide products that are in each sample we can increase the unique identifications in each individual run. We therefore mixed individual samples deriving from different cross-link conditions and analyzed them as a “mixed” sample. Furthermore, optimal cross-linking conditions for the CASP targets were not known to us (and most likely would need to be tailored to each target individually) so given the time constraints of the experiment, it made sense for us to pool different conditions. Typically, mixed samples were injected and technically replicated on LC-MS until subsequent injections failed to yield additional unique residue pairs to the data set (typically three injections) (Supplementary Tables 1-4).

Bottom-up proteomics approaches rely heavily on the near exclusive use of trypsin, with 96% of deposited data having been generated using trypsin (41). Numerous factors have led to this dominance, but essentially reliance on trypsin means that peptide identifications depend on good sequence coverage of tryptic cleavage residues, Arg and Lys. Cross-linking via amine-reactive NHS-ester based cross-linkers (DSS/BS3/sulfo-SDA) targets Lys residues, which subsequently become non-cleavable by trypsin following cross-linking reaction, the prevalence of Lys and Arg residues becomes even more important (42). It has been shown that cross-linked residue pair identification can be boosted by the use of alternative proteases to trypsin, including proteinase K, Asp-N, Glu-C, Lys-C and Lys-N (18, 42, 43). We decided to employ Glu-C to target acidic residues for cleavage for all targets alongside standard trypsin digestion. Trypsin/Glu-C co-digestion was used for Target 1-Tx781 because of the obvious sparseness of tryptic cleavage sites from residues 1-180. The same digestion protocol was used for Target 2-Tx808 but only contributed 15/265 unique residue pairs at 5% FDR (as opposed to 119/305 unique residue pairs at 5% FDR in the case of Target 1-Tx781) so was dropped for remaining targets. For Targets 2-4 (Tx808, Tx767 and Tx812), in-solution Glu-C digestion was used.

The protocol for providing data for CASP11 was identical for each protein target, conforming to a number of steps: (1) buffer-exchange if required; (2) photo-cross-linking, digestion and mass spectrometric analysis; (3) a target-decoy approach FDR analysis and compilation of lists comprising identified residue pairs; (4) submission of our data to CASP11. It should be noted that our data was not posted to the CASP11 participants straight away but subjected to a quality control step by the CASP11 organizers. The CASP11 organizers used their knowledge of the protein structures to verify our confidence intervals (FDR windows) to prevent fundamentally flawed data to be used by the modelers in the second part of the experiment (which wasn’t the case).

We provided the CASP Organizing Committee with linked residue pair lists as .csv files. Three sets of residue pairs were provided in each instance, with 5, 10 and 20% FDR analysis, corresponding to 95%, 90% and 80% residue pair level confidence respectively (Fig. 2). We also provided the CASP Organizing Committee with a Perl script (“CASP_Distances.pl”) along with a ReadMe text file “CASP_Distances_ReadMe.txt” in order to check the distances against the crystal structure held by the organizers. Following validation and without any correction, CASP11 organizers released linked residue pair lists to the registered modelling community. Unbeknown to all participants, the organizers only provided those links that were covered by the crystal structure, which fell short of covering the entire protein in all cases. For the four targets for which cross-linking data was actually acquired, there were 12 and 15 calendar days between data being released to the registered modeling community and the expiration of a target (Fig. 1b).

**FIG. 2.**
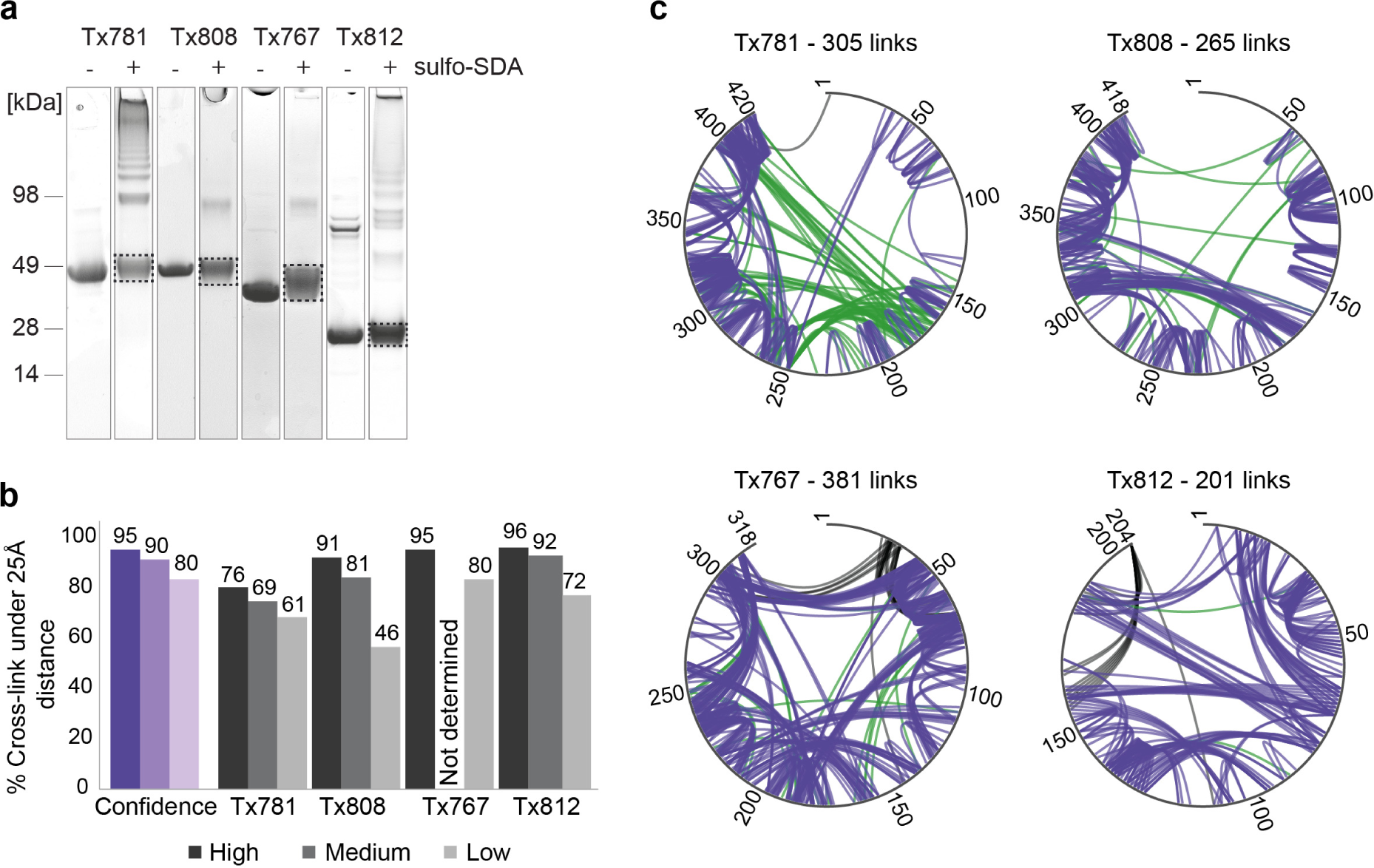
Target cross-linking FDR estimation analysis. (a) CASP11 targets (Tx781, Tx808, Tx767 and Tx812), with (*) and without (-) sulfo-SDA cross-linking. (b) FDR analysis and quality control. FDR estimation on blind data given by *purple columns*. Three groups of data were provided for each target, with differing confidence levels: high (95% true positive hits), medium (90% true positive hits) and low (80% true positive hits). Confidence was inversely related to FDR estimation, which related to FDR cut-offs of 5, 10 and 20%. The black and grey columns represent the results of a data QC check by the CASP Organizing Committee, following submission of cross-linking data by 3DP Lab Edinburgh. Numbers on top of black and grey columns represent the percentage of cross-links found in the known crystal structure that had Cα-Cα cross-linking distances of over 25 Å. (c) Cross-link networks for four CASP targets shown for estimated 5% FDR cut-off. Grey outer lines represent Target sequences. Constraints with Cα-Cα cross-linking distances less than 25 Å are shown in *purple*, constraints with distances 25 Å and over are shown in *green* and constraints missing from the crystal structure and therefore unverifiable are represented in *black*.

### Agreement between CLMS data and solved structures

Results comparisons between CLMS and crystallography are principally challenging as they analyze proteins in different conditions, in solution and in a crystal. As one consequence, crystallography reflects the structure of a single conformer while CLMS samples an ensemble. The methods also return different parameters regarding residue pairs: crystallography returns a distance and CLMS a binary value, seen linked or not, together with a confidence. Here we translate observed links into a 25 Å bound on the distance and hence linkable by photo-CLMS using sulfo-SDA, based on a prior analysis of HSA (28) (in this study, 25 Å is the distance at which the observed Cα-Cα distance distribution merges with the distribution obtained from decoy matches). Note that many links are much shorter than this conservative upper boundary. We assess confidence using a target-decoy approach (Fischer and Rappsilber, submitted). One caveat of our current workflow is that we are missing a site assignment scoring and hence do not control for site assignment errors. Importantly, this does not affect the FDR estimation (decoys model well the distribution of long distance and hence likely false links) and we showed in earlier work that we were able to model HSA domains despite site assignment ambiguities (28), but it may falsely elevate the total number of reported links.

With the exception of the first target, Tx781, the 5% FDR lists matched near perfectly to the crystal structures (Tx808 9%, Tx767 5% and Tx812 4% links >25 Å, respectively, averaging to 6% error) (Figs. 2 and 3). This deteriorated slightly when considering 10% FDR data (19%, n.d., 8%; links >25 Å, respectively, averaging to 10.7% error) and further worsened for 20% FDR data (54%, 20%, 28%; links >25 Å, respectively, averaging to 34% error). We attribute the deviation of the computed to the experimentally assessed accuracy to our still small data sets. FDR using the target-decoy approach relies on large data sets. We can see that this condition is not perfectly fulfilled here. For example, in the case of Tx808 data with 20% FDR added only links with very low score to the 10% FDR list, indicating that the data had been exhaustively matched already at 10% FDR. In conclusion, however, we assess good agreement between photo-CLMS and crystallography in this blind study.

**FIG. 3.**
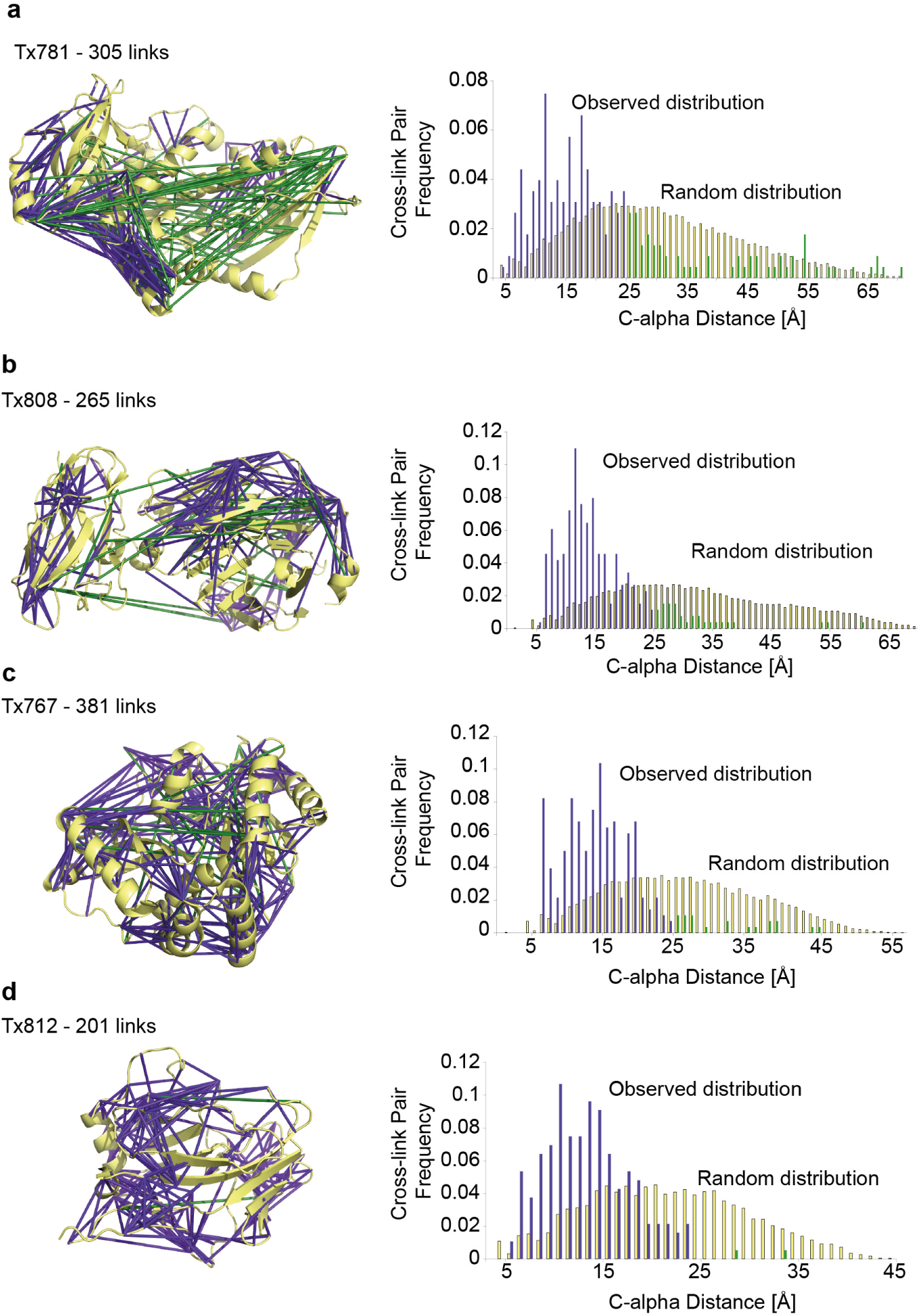
Cross-link distribution within CASP11 Targets. (a) – (d) Left panel shows cross-linked residue pairs at 5% FDR. Right panel shows C-alpha distance distribution of observed constraints at 5% FDR against the random constraint distribution. Constraints with Cα-Cα cross-linking distances less than 25 Å are shown in *purple* and constraints with distances 25 Å and over are shown in *green*. (a) Cross-linked residue pairs of Tx781 in PDB|4qan, n = 305. (b) Cross-linked residue pairs of Tx808 in PDB|4qhw, n = 265. (c) Cross-linked residue pairs of Tx767 in PDB|4qpv, n = 381. (d) Cross-linked residue pairs of Tx812 in crystal structure (*structure not deposited in PDB*), n = 201.

Marked exception from this observation was Tx781. The protein arrived defrosted to the lab and required buffer exchange from Tris to HEPES. Both might have compromised the integrity of the protein structure. Possibly as a result of this or due to cross-linking, the protein was seen as highly aggregated on SDS-PAGE (Fig. 2a). As a further possibility, the protein may have a different structure in solution from in crystal. Our current data does not allow these possibilities to be distinguished.

### Data amount

The number of residue pairs identified in the four proteins using photo-CLMS was comparable to that previously achieved in the analysis of HSA (28) (Table 1). Notably, the acquisition regime used here was more efficient, reducing the acquisition time 3-fold from 12 days to an average of 4.2 days. This is not yet the result of a systematic optimization and further improvements may be possible. Notably, the analysis was not nearing complete detection of linked residue pairs as additional runs kept adding further unique residue pairs, as seen from saturation analysis (Supplementary Figure 1).

**Table 1:** Data amount

| Target protein | MS acquisition time | Links per residue at 5% FDR (number of residue pairs) | Links per residue at 10% FDR (number of residue pairs) | Links per residue at 20% FDR (number of residue pairs) |
| --- | --- | --- | --- | --- |
| HSA (585 AA) | 12 days | 0.85 (500) | 1.51 (881) | 2.56 (1495) |
| Tx781 (420 AA) | 4.7 days | 0.73 (305) | 1.06 (444) | 1.35 (565) |
| Tx808 (418 AA) | 4.4 days | 0.63 (265) | 0.68 (286) | 0.82 (342) |
| Tx767 (318 AA) | 4.0 days | 1.20 (381) | n.d. | 2.26 (718) |
| Tx812 (204 AA) | 3.8 days | 0.99 (201) | 1.30 (265) | 1.77 (360) |

A mitigating factor was that none of the crystal structures covered the entire length of the protein sequence, presumably due to insufficient order in the crystal. Target 3-Tx767 had the highest number of constraints per residue, 1.2 constraints per residue (5% FDR) (Figs. 2c and 3c). 14% of the protein sequence was missing in the crystal structure, however, these missing residues accounted for 27% of identified residue pairs (5% FDR). This reduced the ratio of useful constraints per residue to 0.88 for our experiment.

In the current way cross-links are being used to assist protein structure prediction, special value is given to residue pairs that are far apart in the primary sequence of the protein. Residue pairs close in sequence only provide information relating to local secondary structure, which is less informative for modelling. However, residue pairs that are far apart (>11 residues) in sequence capture information about the spatial arrangement of secondary structures, and therefore a cut-off (>11 sequence separation only) is often used in modelling. Belsom *at al*. used only links larger than 11 residues apart in their modelling of HSA domains (28). Also in this respect, the current data is comparable to the previous study on HSA (Table 2).

**Table 2:** Number of identified residue pairs >11 residues apart in protein sequence

| Target protein | 5% FDR | 5-10% FDR | 10-20% FDR |
| --- | --- | --- | --- |
| HSA (585 AA) | 330 (66%) | 292 (77%) | 511 (83%) |
| Tx781 (420 AA) | 189 (62%) | 100 (72%) | 99 (82%) |
| Tx808 (418 AA) | 155 (59%) | 15 (71%) | 52 (93%) |
| Tx767 (318 AA) | 277 (73%) | n.d. | 234 (69%) |
| Tx812 (204 AA) | 146 (73%) | 48 (75%) | 63 (66%) |

### Protein structure modeling in CASP11 with CLMS data

In this section, we briefly summarize the modelling results in CASP using CLMS data. A full report from the protein modelling perspective is published elsewhere (36). Protein targets accompanied by CLMS data are difficult to model and either lack templates in the PDB or the template structures cannot be found using current template identification methods. These proteins are called free modelling (FM targets) in CASP11. Thus, FM targets test the ability of *ab initio* structure prediction using CLMS data. Note that the prediction groups are not required to use *ab initio* methods for proteins of this category. Therefore, predictors might use *ab initio* structure prediction, remote homology modeling, or hybrid algorithms that combine the two approaches to model these proteins. However, this experiment comes closest to a realistic *ab initio* structure prediction scenario.

Cross-links effectively provide distance restraints between residue pairs with an upper distance bound (25 Å between carbon atoms in this case). This information is usually used to add a restraint term to the energy function used in protein modelling. This restraint term measures the degree of which a given protein conformation satisfies the cross-link restraints and therefore guides structure sampling towards protein conformations that agree with the experimental data.

We compared the performance of CLMS assisted predictions and unassisted predictions from groups that participated in both experiments. We assume that these groups use similar algorithms to perform predictions with and without CLMS data. Therefore, comparing these groups should capture the impact of CLMS data. For these 19 groups, the CLMS data slightly improved the GDT_TS of the predicted models from 36.4 to 38.1 for the first and from 40.9 to 42.0 for the best-of-five submitted models. The GDT_TS is a measure for the match of the prediction to the native structure and ranges from 0 (structures completely dissimilar) to 100 (perfect match).

Some groups submitted superior CLMS assisted predictions but correspondence with these groups suggests that CLMS data was not the driving factor behind these predictions. Furthermore, when we consider all predictions in CASP11, the best models from the unassisted category are still better than CLMS assisted models (mean GDT_TS of 40 and 37.5, respectively). One possible explanation for this finding is that many more prediction groups (143) submitted models to the unassisted category then to the CLMS assisted category (19). Overall, the results suggest a slight increase in CLMS-assisted modelling accuracy when comparing groups that participated in both experiments but there is no clear, overall improvement of CLMS assisted predictions. However, the CLMS-CASP experiment was helpful to identify experimental challenges for current CLMS protocols in structure modelling, which we will discuss in the next section.

### Sequence coverage

The observed cross-links did not evenly distribute throughout the entire structure of the targets. This issue was particularly prevalent for Target 1-Tx781 (Fig. 4a). There was a notable absence of cross-linked residue pairs involving residues up to residue 180 (0.11 distance constraints per residue), compared with the number observed involving residue 181 onwards (0.69 distance constraints per residue). The absence of observed residue pairs correlates with low frequency of tryptic cleavage sites: up to residue 180 there are 18 tryptic cleavage sites, compared with 31 in the remaining 224 residues. Furthermore, if lysine residues are significantly decorated by sulfo-SDA, the extent of proteolytic cleavage by trypsin becomes more dependent on the availability of arginine residues. There are 10 arginine residues in the second part of the sequence compared to only 3 in the sequence up to residue 180. The sequence distribution of lysine and arginine residues is also an important factor, which dictates whether observation of residue pairs is likely or not. If K126 of Tx781 is modified or cross-linked the resulting tryptic peptide would be 60 residues long; prohibitively long for ordinary mass spectrometry analysis. We would not realistically expect to observe residue pairs involving residues in this stretch of the sequence (14% of the total sequence). Likewise, modified or cross-linked K93 and K47 would create 48- and 36-residue tryptic peptides, respectively. A total of 111 residues (62% of the sequence up to residue 180, 26% of the total protein sequence) are thus theoretically inaccessible via trypsin digestion. Conversely, residue pairs involving K154 and K156 would be inaccessible because the resulting tryptic peptides would be too short, at 3 and 4 amino acids long respectively, as the minimum peptide length for the analysis was stipulated as 5 residues.

**FIG. 4.**
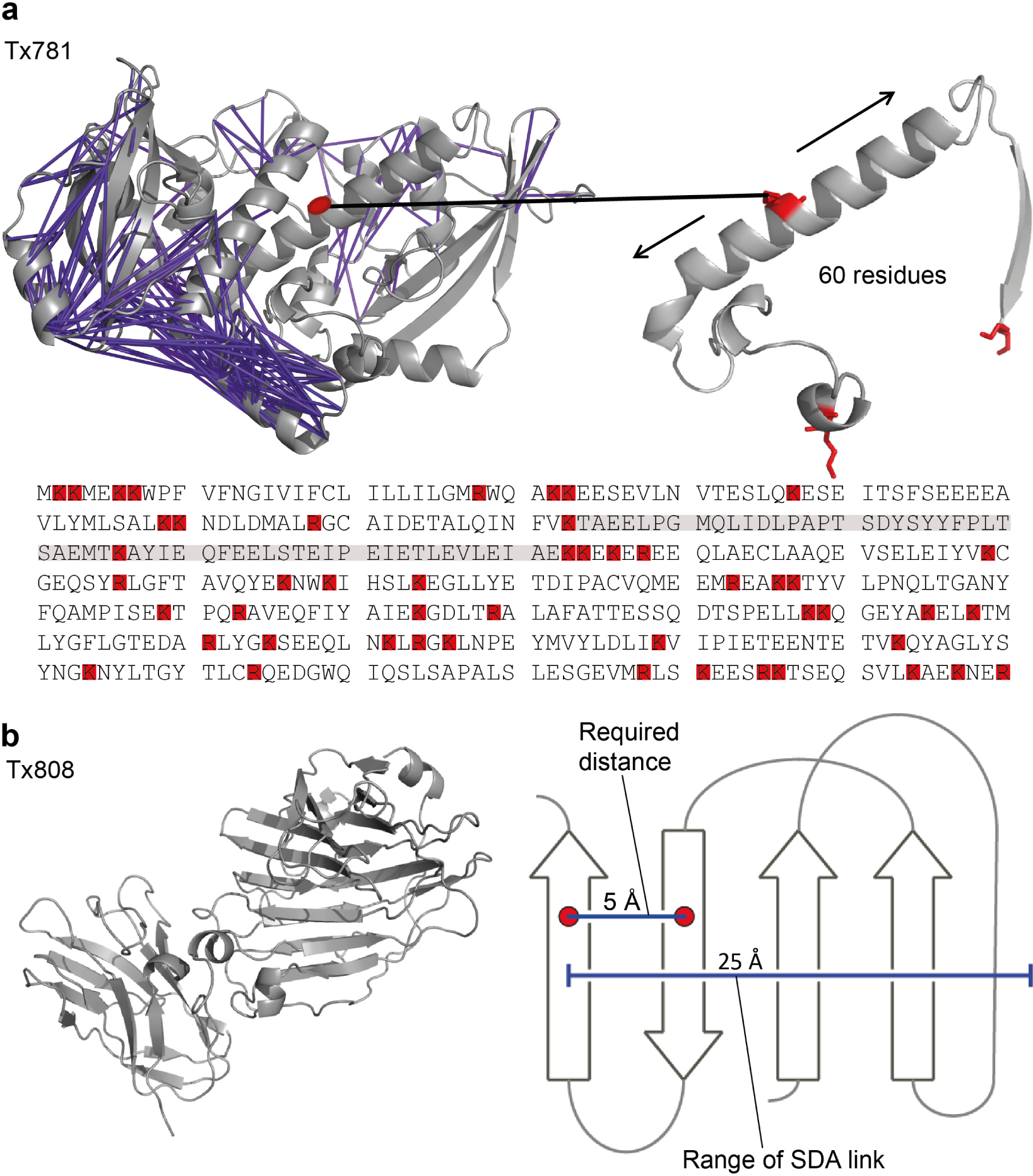
Procedural limitations identified in the study. (a) *Top left*: Constraints under 25 Å shown in *purple*, in the crystal structure of Tx781 (PDB|4qan). *Top right*: Zoom of a 61 amino acid tryptic peptide devoid of observed constraints, containing a single, centrally located lysine residue highlighted in *red*. *Bottom*: Amino acid sequence of Tx781. Tryptic (lys and arg) residues highlighted in *red*. (b) *Left*: Tx808 crystal structure (PDB|4qhw). *Right*: Representation of a beta-sheet.

If our reasoning were correct one would expect digestion with alternative enzymes to lead to improved coverage, especially in the N-terminal region of Tx781. This was indeed the case, as alternative digestion strategies including Glu-C, rather than relying on digestion by trypsin alone yielded substantial improvements (Table 3). Glutamic acid residues are frequent in the protein (61 in total (15% of total residues) and 32 in residues 1 to 180 of the sequence (18% of residues)). Digestion methods including Glu-C contributed 39% of the total residue pairs identified at 5% FDR for Tx781. In the N-terminal region of Tx781, the proportion of residue pairs identified through addition of Glu-C digestion rose to 74%, with data amount was increased from 0.11 to 0.43 distance constraints per residue (at 5% FDR), compared with using trypsin digestion alone. For residue pairs starting at residue 181, double-digestion increased data amount from 0.69 to 0.95 distance constraints per residue (at 5% FDR), compared with using trypsin digestion alone. Use of double-digestion strategies had a higher impact in the parts of the protein lacking tryptic cleavage sites, as was initially expected.

**Table 3:**
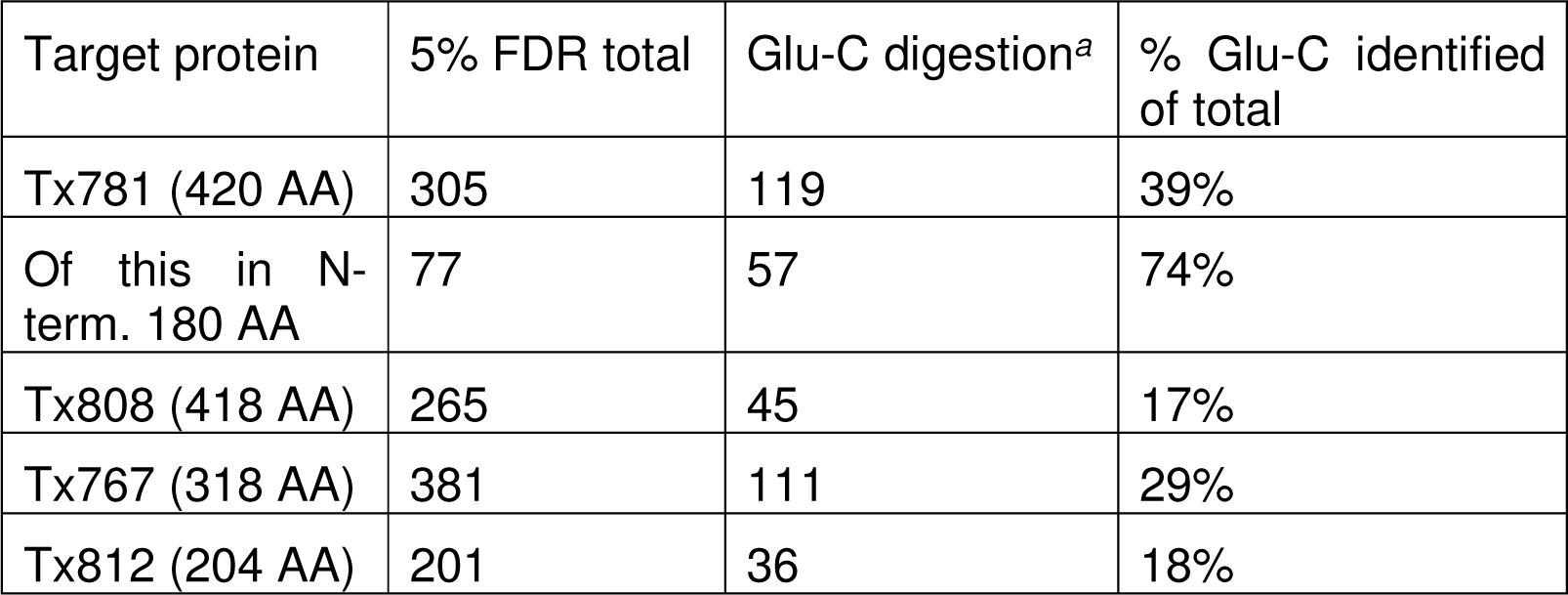
Number of identified residue pairs added (at 5% FDR) with alternatives to trypsin-only digestion. *^a^* All digestion methods involving use of Glu-C (including trypsin/Glu-C co-digestion (Tx781 and Tx808), and in-solution Glu-C digestion (Tx808, Tx767 and Tx812).

More modest gains were made with Tx808, which was due in part to a lack of both tryptic and Glu-C cleavage sites in the protein, and with Tx767 and Tx812, where a lack of tryptic cleavage sites was less of a concern.

### Beta sheet issues

Beta sheet topology is present in all targets with Target 2-Tx808 being the extreme; 54% of residues (215 of 400 residues present in the x-ray crystal structure) are in beta strands (Figs. 4b and 5a). We noticed a marked bias against cross-links in the beta sheet region of the protein (Fig. 5b). Cross-links are found predominantly in the loop regions of the protein (Fig. 3b). For Tx808, 64% of observed residue pairs (170 of 265, at 5% FDR) involve at least one beta strand residue in the pair. However, if there were no bias against cross-linked residue pairs in beta strands, one would expect this number to be 80%. Furthermore, if there were no bias against cross-linked residue pairs in beta sheets, one would expect 30% of residue pairs where both residues have beta strand structure. We observe this to be only 21% (Fig. 5b). The same discrepancy between the expected numbers of residue pairs identified with both residues having beta strand structure and the actual number identified in the datasets is observed for Tx767 and Tx812 (we exclude Tx781 from our analysis because of the issues discussed earlier), as seen in Fig. 5b. Tx767 has the lowest percentage of residues in the crystal structure having beta strand structure, so hence one would expect fewer residue pairs between beta strand residues (3.8% of all residue pairs), however only 0.30% of identified residue pairs in our 5% FDR dataset match this condition.

**FIG. 5.**
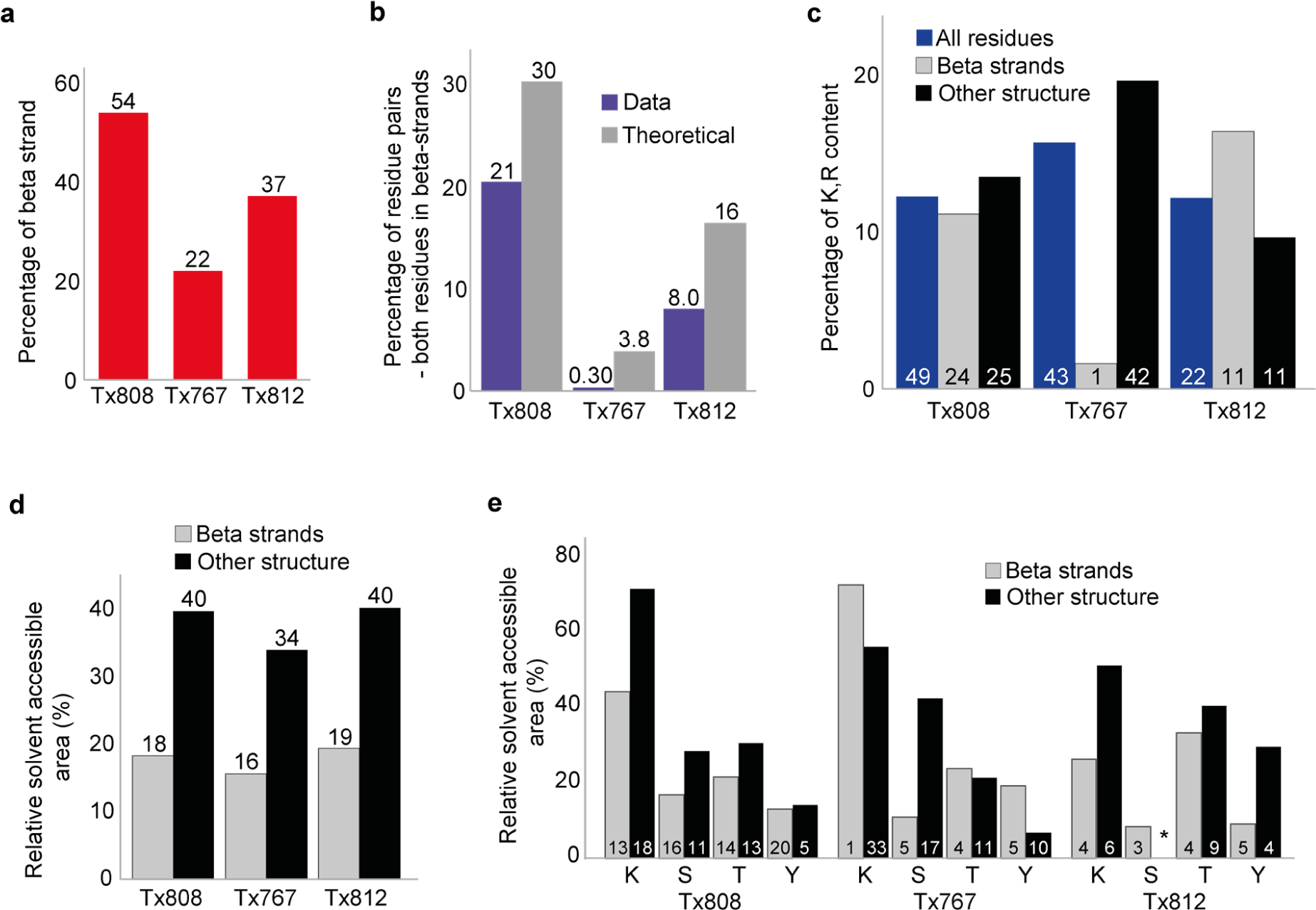
Beta strand analysis for targets Tx808, Tx767, Tx812. (a) Percentage of residues with beta strand structure. (b) Percentage of residue pairs where both residues are beta strand in identified cross-link data compared with the dataset where all possible residue pairs are considered. Residue pairs identified at 5% FDR shown by *purple columns*. All possible residue pairs with both residues in beta strand shown by *grey columns*. (c) Percentage of Lys and Arg residues in beta strands versus residues with all other structure types. Percentages of Lys and Arg residues across all residues for each target are shown by *blue columns*. Percentages of Lys and Arg residues in beta strands for each target are shown by *grey columns*. Percentage of Lys and Arg residues for all residues, excluding those with beta strand structure are shown by *black columns*. Numbers within columns refer to the numbers of Lys and Arg found in the crystal structure of each target. (d) Average relative solvent accessible area (RSA) of residues for each target in beta strands compared with average RSA for all other residues, excluding beta strand residues. Average RSAs of beta strand residues are shown by *grey columns*. Average RSAs of all other residues, excluding beta strand residues, are shown by *black columns*. (e) Average RSA of Lys, Ser, Thr and Tyr residues for each target, with RSA values of these residues in beta strands compared with values for residues in all other residues, excluding beta strand residues. Beta strand Lys, Ser, Thr and Tyr are shown by *grey columns*. Lys, Ser, Thr and Tyr from all other residues, excluding beta strand residues, are shown by *black columns*. Numbers within columns refer to the number of each type of residue found in the crystal structure of each target.

We hypothesized that a lack of tryptic cleavage sites in beta strands might lead to an absence of cross-linked beta strands. We looked at the percentage of lysine and arginine residues in all residues with beta strand structure for each target, in comparison with the percentage of lysine and arginine residues in other structure types, as well as the percentage of lysine and arginine over all residues found in the crystal structures (Fig. 5c). For Tx767, this hypothesis could indeed be correct. There is only 1 lysine residue with beta strand structure in Tx767 and no arginine residues. This not only affects the digestability of this region but also the possibility of anchoring the reagent. Tx808 and Tx812 have a larger percentage of tryptic residues in beta strands. Consequently, digestion should be less of a concern here. However, these proteins also have a lower percentage of residues that react with NHS-esters in beta-strands than other structure elements. While lack of trypsin cleavage sites may account for the problems encountered in the beta sheets of Tx767 they do not account for the problems encountered in the beta sheets of Tx808 and Tx812. Yet, availability of anchoring residues may be a general concern.

We next considered relative solvent accessibility (RSA) (44, 45) and whether this could explain discrepancies in expected and observed cross-link patterns. On average, beta strand resides have much lower relative solvent accessible areas in each of the targets (16-19%) as opposed to other residues (34-40%) (Fig. 5d). This means that on average, residues contained within beta strands are more buried and solvent inaccessible than residues with other structure. The same disparity between beta strand residues and other residues holds true when looking at the NHS-ester reactive residues, Lys, Ser, Thr and Tyr (Fig. 5e). These residues must be sufficiently accessible for the first anchoring step of cross-linking with sulfo-SDA to occur. Note that there is only a single Lys residue in beta strands of Tx767, which happens to have a high RSA and additionally, Thr residues have a moderately higher RSA for this protein. Consequentially, both lack of anchoring residues, cleavage sites and lower RSA may contribute in different ratios and in different proteins to the problem of beta sheet analysis by CLMS.

There is an even more fundamental further issue concerning the current approach towards beta-sheet topology. This has its origins in the use of sulfo-SDA as cross-linker (Fig. 4b). The alpha-carbon distance between adjacent hydrogen bonded beta strands is in the order of 5 Å (46), however the upper limit distance boundary defined by SDA is in the range of 20 to 25 Å. This covers as much as five beta strands. Such a spatial resolution especially at current data density is insufficient to reveal the topology of beta strands in a beta sheet.

## CONCLUSIONS

Our test of CLMS under the auspices of CASP11 exposed pitfalls and problems that are likely to be encountered for a large proportion of proteins with currently unsolved structure. While SDA proved very useful in solving the structure of the alpha-helical domains of HSA (28) CLMS at large may require major developments to achieve a similar success for beta-sheet proteins. Sequence coverage and spatial resolution pose a technological challenge and spell out the agenda for future developments. We could show that a blind study to test CLMS can be organized and is successful in revealing current weaknesses of at least the workflow employed by us and for single protein analysis. Multi-protein complexes or large protein networks may pose additional challenges that remain invisible by this analysis. To reveal those it would be desirable to develop complementary tests, albeit simply extending the current set-up will be difficult. The number of large multi-protein complexes to see their structure solved is small and so is the chance of coordinating their public release with a blind CLMS study. In any case, CASP organizers indicated their willingness to open also their next round starting in May 2016, to the CLMS community. We would like to extend an open call to any interested lab to seize this opportunity to thoroughly and openly test their CLMS workflows with regards to single protein analysis.

## Acknowledgments

We would like to thank the CASP11 Organizing Committee, in particular Krzysztof Fidelis and Andriy Kryshtafovych for their coordination efforts, PDB for their cooperation in delaying the release of one target by one week and the following researchers for providing protein targets: Mark Wilson and Janani Prahlad (Department of Biochemistry/Redox Biology, University of Nebraska), Ashley M. Deacon and Qingping Xu (Joint Center for Structural Genomics (JCSG), Stanford Synchrotron Radiation Lightsource, Stanford University), Gaetano Montelione and Rong Xiao (Center for Advanced Biotechnology and Medicine, Rutgers University), Jörg Martin (Max-Planck Institute for Developmental Biology, Tübingen), Deborah Fass (Department of Structural Biology, Weizmann Institute of Science). We acknowledge the PRIDE team for the deposition of our data to the ProteomeXchange Consortium. The MS data have been deposited to the ProteomeXchangeConsortium (http://proteomecentral.proteomexchange.org) via the PRIDE partner repository with the dataset identifier PXD003643 (40). This work was generously supported by the Wellcome Trust (Senior Research Fellowship to JR 103139, instrument grant 108504 and Centre core grant 092076), by the Alexander-von-Humboldt foundation through funding from the German Federal Ministry of Education and Research (BMBF) and by NIH Grant (1 R01 GM076706).

## REFERENCES

1. Rappsilber, J. (2011) The beginning of a beautiful friendship: cross-linking/mass spectrometry and modelling of proteins and multi-protein complexes. J Struct Biol 173, 530–540

2. Paramelle, D., Miralles, G., Subra, G., and Martinez, J. (2013) Chemical cross-linkers for protein structure studies by mass spectrometry. Proteomics 13, 438–456

3. Walzthoeni, T., Leitner, A., Stengel, F., and Aebersold, R. (2013) Mass spectrometry supported determination of protein complex structure. Curr Opin Struct Biol 23, 252–260

4. Sinz, A. (2006) Chemical cross-linking and mass spectrometry to map three-dimensional protein structures and protein-protein interactions. Mass Spectrometry Reviews 25, 663–682

5. Leitner, A., Faini, M., Stengel, F., and Aebersold, R. (2016) Crosslinking and Mass Spectrometry: An Integrated Technology to Understand the Structure and Function of Molecular Machines. Trends Biochem Sci 41, 20–32

6. Rappsilber, J., Siniossoglou, S., Hurt, E. C., and Mann, M. (2000) A generic strategy to analyze the spatial organization of multi-protein complexes by cross-linking and mass spectrometry. Anal Chem 72, 267–275

7. Maiolica, A., Cittaro, D., Borsotti, D., Sennels, L., Ciferri, C., Tarricone, C., Musacchio, A., and Rappsilber, J. (2007) Structural analysis of multiprotein complexes by cross-linking, mass spectrometry, and database searching. Mol Cell Proteomics 6, 2200–2211

8. Chen, Z. A., Jawhari, A., Fischer, L., Buchen, C., Tahir, S., Kamenski, T., Rasmussen, M., Lariviere, L., Bukowski -Wills, J. C., Nilges, M., Cramer, P., and Rappsilber, J. (2010) Architecture of the RNA polymerase II–TFIIF complex revealed by cross -linking and mass spectrometry. EMBO J 29, 717–726

9. Lauber, M. A., Rappsilber, J., and Reilly, J. P. (2012) Dynamics of ribosomal protein S1 on a bacterial ribosome with cross-linking and mass spectrometry. Mol Cell Proteomics 11, 1965–1976

10. Kim, S. J., Fernandez-Martinez, J., Sampathkumar, P., Martel, A., Matsui, T., Tsuruta, H., Weiss, T. M., Shi, Y., Markina-Inarrairaegui, A., Bonanno, J. B., Sauder, J. M., Burley, S. K., Chait, B. T., Almo, S. C., Rout, M. P., and Sali, A. (2014) Integrative structure-function mapping of the nucleoporin Nup133 suggests a conserved mechanism for membrane anchoring of the nuclear pore complex. Mol Cell Proteomics 13, 2911–2926

11. Shi, Y., Fernandez-Martinez, J., Tjioe, E., Pellarin, R., Kim, S. J., Williams, R., Schneidman-Duhovny, D., Sali, A., Rout, M. P., and Chait, B. T. (2014) Structural characterization by cross-linking reveals the detailed architecture of a coatomer-related heptameric module from the nuclear pore complex. Mol Cell Proteomics 13, 2927–2943

12. Gordiyenko, Y., Schmidt, C., Jennings, M. D., Matak-Vinkovic, D., Pavitt, G. D., and Robinson, C. V. (2014) eIF2B is a decameric guanine nucleotide exchange factor with a γ2ε2 tetrameric core. Nat Commun 5, 3902

13. Greber, B. J., Boehringer, D., Leibundgut, M., Bieri, P., Leitner, A., Schmitz, N., Aebersold, R., and Ban, N. (2014) The complete structure of the large subunit of the mammalian mitochondrial ribosome. Nature 515, 283–U326

14. Greber, B. J., Bieri, P., Leibundgut, M., Leitner, A., Aebersold, R., Boehringer, D., and Ban, N. (2015) The complete structure of the 55S mammalian mitochondrial ribosome. Science 348, 303–308

15. Tomko, R. J., Taylor, D. W., Chen, Z. A., Wang, H. W., Rappsilber, J., and Hochstrasser, M. (2015) A Single α Helix Drives Extensive Remodeling of the Proteasome Lid and Completion of Regulatory Particle Assembly. Cell 163, 432–444

16. Barysz, H., Kim, J. H., Chen, Z. A., Hudson, D. F., Rappsilber, J., Gerloff, D. L., and Earnshaw, W. C. (2015) Three-dimensional topology of the SMC2/SMC4 subcomplex from chicken condensin I revealed by cross-linking and molecular modelling. Open Biol 5, 150005

17. Fischer, L., Chen, Z. A., and Rappsilber, J. (2013) Quantitative cross-linking/mass spectrometry using isotope-labelled cross-linkers. J Proteomics 88, 120–128

18. Leitner, A., Reischl, R., Walzthoeni, T., Herzog, F., Bohn, S., Förster, F., and Aebersold, R. (2012) Expanding the chemical cross-linking toolbox by the use of multiple proteases and enrichment by size exclusion chromatography. Mol Cell Proteomics 11, M111.014126

19. Young, M. M., Tang, N., Hempel, J. C., Oshiro, C. M., Taylor, E. W., Kuntz, I. D., Gibson, B. W., and Dollinger, G. (2000) High throughput protein fold identification by using experimental constraints derived from intramolecular cross-links and mass spectrometry. Proc Natl Acad Sci U S A 97, 5802–5806

20. Kwan, A. H., Mobli, M., Gooley, P. R., King, G. F., and Mackay, J. P. (2011) Macromolecular NMR spectroscopy for the non-spectroscopist. FEBS J 278, 687–703

21. Gomes, A. F., and Gozzo, F. C. (2010) Chemical cross-linking with a diazirine photoactivatable cross-linker investigated by MALDI- and ESI-MS/MS. J Mass Spectrom 45, 892–899

22. Piotrowski, C., Ihling, C. H., and Sinz, A. (2015) Extending the cross-linking/mass spectrometry strategy: Facile incorporation of photo-activatable amino acids into the model protein calmodulin in Escherichia coli cells. Methods 89, 121–127

23. Suchanek, M., Radzikowska, A., and Thiele, C. (2005) Photo-leucine and photo-methionine allow identification of protein-protein interactions in living cells. Nat Methods 2, 261–267

24. Ječmen, T., Ptáčková, R., Černá, V., Dračínská, H., Hodek, P., Stiborová, M., Hudeček, J., and Šulc, M. (2015) Photo-initiated crosslinking extends mapping of the protein-protein interface to membrane-embedded portions of cytochromes P450 2B4 and b5. Methods 89, 128–137

25. Hétu, P. O., Ouellet, M., Falgueyret, J. P., Ramachandran, C., Robichaud, J., Zamboni, R., and Riendeau, D. (2008) Photo-crosslinking of proteins in intact cells reveals a dimeric structure of cyclooxygenase-2 and an inhibitor-sensitive oligomeric structure of microsomal prostaglandin E2 synthase-1. Arch Biochem Biophys 477, 155–162

26. Kolbel, K., Ihling, C. H., and Sinz, A. (2012) Analysis of peptide secondary structures by photoactivatable amino acid analogues. Angew Chem Int Ed Engl 51, 12602–12605

27. Lössl, P., Kölbel, K., Tänzler, D., Nannemann, D., Ihling, C. H., Keller, M. V., Schneider, M., Zaucke, F., Meiler, J., and Sinz, A. (2014) Analysis of nidogen-1/laminin γ1 interaction by cross-linking, mass spectrometry, and computational modeling reveals multiple binding modes. PLoS One 9, e112886

28. Belsom, A., Schneider, M., Fischer, L., Brock, O., and Rappsilber, J. (2016) Serum Albumin Domain Structures in Human Blood Serum by Mass Spectrometry and Computational Biology. Mol Cell Proteomics 15, 1105–1116

29. Walzthoeni, T., Claassen, M., Leitner, A., Herzog, F., Bohn, S., Forster, F., Beck, M., and Aebersold, R. (2012) False discovery rate estimation for cross-linked peptides identified by mass spectrometry. Nature Methods 9, 901–903

30. Brunette, T. J., and Brock, O. (2005) Improving protein structure prediction with model-based search. Bioinformatics 21 Suppl 1, i66–74

31. Moult, J., Pedersen, J. T., Judson, R., and Fidelis, K. (1995) A large-scale experiment to assess protein structure prediction methods. Proteins 23, ii–v

32. Kryshtafovych, A., Monastyrskyy, B., and Fidelis, K. (2016) CASP11 statistics and the prediction center evaluation system. Proteins

33. Moult, J. (2005) A decade of CASP: progress, bottlenecks and prognosis in protein structure prediction. Curr Opin Struct Biol 15, 285–289

34. Moult, J., Fidelis, K., Kryshtafovych, A., Schwede, T., and Tramontano, A. (2014) Critical assessment of methods of protein structure prediction (CASP)–round x. Proteins 82 Suppl 2, 1–6

35. Kryshtafovych, A., Moult, J., Bales, P., Bazan, J. F., Biasini, M., Burgin, A., Chen, C., Cochran, F. V., Craig, T. K., Das, R., Fass, D., Garcia-Doval, C., Herzberg, O., Lorimer, D., Luecke, H., Ma, X., Nelson, D. C., van Raaij, M. J., Rohwer, F., Segall, A., Seguritan, V., Zeth, K., and Schwede, T. (2014) Challenging the state of the art in protein structure prediction: Highlights of experimental target structures for the 10th Critical Assessment of Techniques for Protein Structure Prediction Experiment CASP10. Proteins 82 Suppl 2, 26–42

36. Schneider, M., Belsom, A., Rappsilber, J., and Brock, O. (2016) Blind Testing of Cross-linking/Mass Spectrometry Hybrid Methods in CASP11. Proteins

37. Rappsilber, J., Ishihama, Y., and Mann, M. (2002) Stop and Go Extraction Tips for Matrix-Assisted Laser Desorption/Ionization, Nanoelectrospray, and LC/MS Sample Pretreatment in Proteomics. Analytical Chemistry 75, 663–670

38. Ishihama, Y., Rappsilber, J., Andersen, J. S., and Mann, M. (2002) Microcolumns with self-assembled particle frits for proteomics. Journal of Chromatography A 979, 233–239

39. Cox, J., and Mann, M. (2008) MaxQuant enables high peptide identification rates, individualized p.p.b.-range mass accuracies and proteome-wide protein quantification. Nat Biotechnol 26, 1367–1372

40. Vizcaíno, J. A., Deutsch, E. W., Wang, R., Csordas, A., Reisinger, F., Ríos, D., Dianes, J. A., Sun, Z., Farrah, T., Bandeira, N., Binz, P. A., Xenarios, I., Eisenacher, M., Mayer, G., Gatto, L., Campos, A., Chalkley, R. J., Kraus, H. J., Albar, J. P., Martinez-Bartolomé, S., Apweiler, R., Omenn, G. S., Martens, L., Jones, A. R., and Hermjakob, H. (2014) ProteomeXchange provides globally coordinated proteomics data submission and dissemination. Nat Biotechnol 32, 223–226

41. Tsiatsiani, L., and Heck, A. J. (2015) Proteomics beyond trypsin. FEBS J 282, 2612–2626

42. Petrotchenko, E. V., Serpa, J. J., Hardie, D. B., Berjanskii, M., Suriyamongkol, B. P., Wishart, D. S., and Borchers, C. H. (2012) Use of proteinase K nonspecific digestion for selective and comprehensive identification of interpeptide cross-links: application to prion proteins. Mol Cell Proteomics 11, M111.013524

43. Leitner, A., Joachimiak, L. A., Unverdorben, P., Walzthoeni, T., Frydman, J., Förster, F., and Aebersold, R. (2014) Chemical cross-linking/mass spectrometry targeting acidic residues in proteins and protein complexes. Proc Natl Acad Sci U S A 111, 9455–9460

44. Lee, B., and Richards, F. M. (1971) The interpretation of protein structures: estimation of static accessibility. J Mol Biol 55, 379–400

45. Chothia, C. (1976) The nature of the accessible and buried surfaces in proteins. J Mol Biol 105, 1–12

46. Levitt, M., and Chothia, C. (1976) Structural patterns in globular proteins. Nature 261, 552–558

